# Neurometabolic signaling and control of policy complexity

**DOI:** 10.1101/2025.02.24.639890

**Authors:** Alexander L. Tesmer, Christine Dalla Pola, Dino Gilli, Nikola Grujic, Eva F. Bracey, Tommaso Patriarchi, Daria Peleg-Raibstein, Rafael Polania, Denis Burdakov

**Affiliations:** Swiss Federal Institute of Technology (ETH Zürich), Department of Health Sciences and Technology, Zürich, Switzerland; Neuroscience Center Zürich (ZNZ); Institute of Pharmacology and Toxicology, University of Zürich, Zürich, Switzerland

## Abstract

Cognition and adaptive behavior emerge from neural information processing. This must operate within finite metabolic constraints, since neural information processing is metabolically expensive. While neural implementations of action selection and learning are well-studied, systems allocating the informational capacity required to encode complex behavioral policies remain unknown. We hypothesized that hypothalamic hypocretin/orexin neurons (HONs) are uniquely positioned to signal and control policy complexity, given that they are activated by metabolic depletion and influence decision-making systems. To explore this, we employed a set of cell/neurotransmitter-specific imaging and causal manipulations during a multi-armed bandit task where freely behaving mice learned probabilistic state-action-reward relationships (together ∼100,000 decisions from >100 mice). Miniscope recordings of HON activity revealed that pre-choice, but not post-choice activity correlates with decision policy, dissociating decisions from feedback. Furthermore, manipulating HON signals with optogenetics and pharmacology confirmed that they causally regulate the development of complex policies. Finally, neurotransmitter-specific sensors revealed that hypocretin/orexin receptors modulate decision policy-related dopamine and noradrenaline dynamics in the nucleus accumbens and medial prefrontal cortex. These findings identify HONs as subcortical regulators of policy complexity, encoding critical signals for decision-making adjustments. This opens a new window for the development of comprehensive mechanistic models of strategic learning which account for interplay between decision-making policies and metabolic/informational constraints, and may guide the development of potential treatments for disorders with policy deficits such as autism and schizophrenia.

## INTRODUCTION

Strategic learning in systems with metabolic or informational constraints relies on two key components: (i) the set of *learning rules* applied based on action, feedback, and experiences with the environment, and (ii) the *decision policies* determining not only what decision to make but also how much of the limited metabolic or informational budget to allocate to each decision instance. While a significant portion of the literature has focused on understanding learning rules and related processes through the lens of, for instance, dopaminergic systems, the neural mechanisms governing the allocation of limited metabolic or informational resources during decision-making remain poorly understood.

In recent years, there has been growing interdisciplinary interest in neuroscience^1, 2^, psychology^3–5^, economics^6–8 9^, and machine learning^10, 11^ to understand how agents—whether biological or artificial—learn to interact with uncertain environments to maximize reward- or fitness-related objective functions under resource constraints. Many of these advancements are rooted in foundational principles from information theory^12–15^, and empirical work on sensory systems ^16^ which has demonstrated the role of metabolic and energetic constraints in shaping information processing^17^, revealing how biological systems optimize resource use while maximizing informational or behavioral objectives^18–20 21, 22^. Recent work has expanded this understanding beyond the sensory domain by studying how policy complexity and informational constraints influence decision-making when learning to navigate environments under uncertainty^23, 24^. Crucially, this line of work emphasizes that agents often face a trade-oI between the complexity of their decision policies and the computational or metabolic costs of implementing them^25, 26^. While empirical work and computational models appear to support the idea that our behavior approximates optimal policy compression under resource limitations, the neural mechanisms by which the brain signals and controls the complexity of behavioral policies remain poorly understood. Bridging this gap may thus require a tight integration of empirical neuroscience with computational frameworks to better understand the interplay between neural dynamics, environmental complexity, and adaptive behavior.

Complexity, quantified in bits, is metabolically expensive; it costs approximately 10^4^ ATP molecules to transmit one bit of information at a chemical synapse ^17, 27, 28^. We hypothesized that neural systems tracking metabolic states and influencing decision-making systems are well-poised to regulate and promote the allocation of complexity in behavioral policies; governing the implementation of strategic decision policies while considering, for instance, the metabolic state of the organism. One such candidate complying with these requirements is the system instantiated by hypothalamic hypocretin/orexin-producing neurons (HONs) ^29–33^. HONs track the slowly changing metabolic states^29, 34–45^, but also generate more rapidly-varying and less understood phasic signals^41, 46–51^; regulate arousal^52–56^; and send outputs to multiple downstream neural systems implicated in decisions and learning, including dopaminergic and noradrenergic systems^57–64^.

## RESULTS

### Quantifying behavioral policy in mice

To test this hypothesis, we implemented a multiarmed bandit task allowing us to examine the probabilistic relationships between actions, rewards, and states. We placed n = 97 BL/6J mice (n = 42 male and n = 55 female) into a 3-arm Y-Maze where each equidistant arm contained a lick-activated port distributing (probabilistic) sucrose reward (Fig. 1A, see methods). Each session consisted of two phases: a 5 minutes deterministic phase, serving as an adaptation period, in which reward probability was always 100%, and then a 25 minutes probabilistic phase where each port was pseudo-randomly assigned reward probability 25%, 50%, or 100%. Critically, a reward could not be dispensed from the same port twice in a row, thus creating three unique states *s* for the task. In each state, mice had to implement an action *a* indicating a *left* port or *right* decision. For example, in state 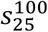 mice can only choose between the 100% and 25% probability port after the 50% port was selected in the previous trial (Fig. B). In each session mice continuously performed trials resulting in an average of 131 ± 4 trials across mice (Fig. 1C). Mice learned to choose the 100% port and were less likely to choose the 25% port (rmANOVA *P*<0.001, Fig. 1D-E) and the expected reward in the probabilistic phase (60.9% ± 0.3%) was better than chance (P<0.001, Fig. 1F), suggesting mice learned the task contingencies. Finally, we found performance was highest in state 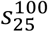, while states 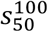 and 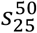 proved more difficult, indicating task performance was state-dependent (rmANOVA *P*<0.001, Fig. 1G). Broadly, we found minimal sex differences, although males tended to perform slightly more trials per session (Fig. S1).

**Fig. 1.**
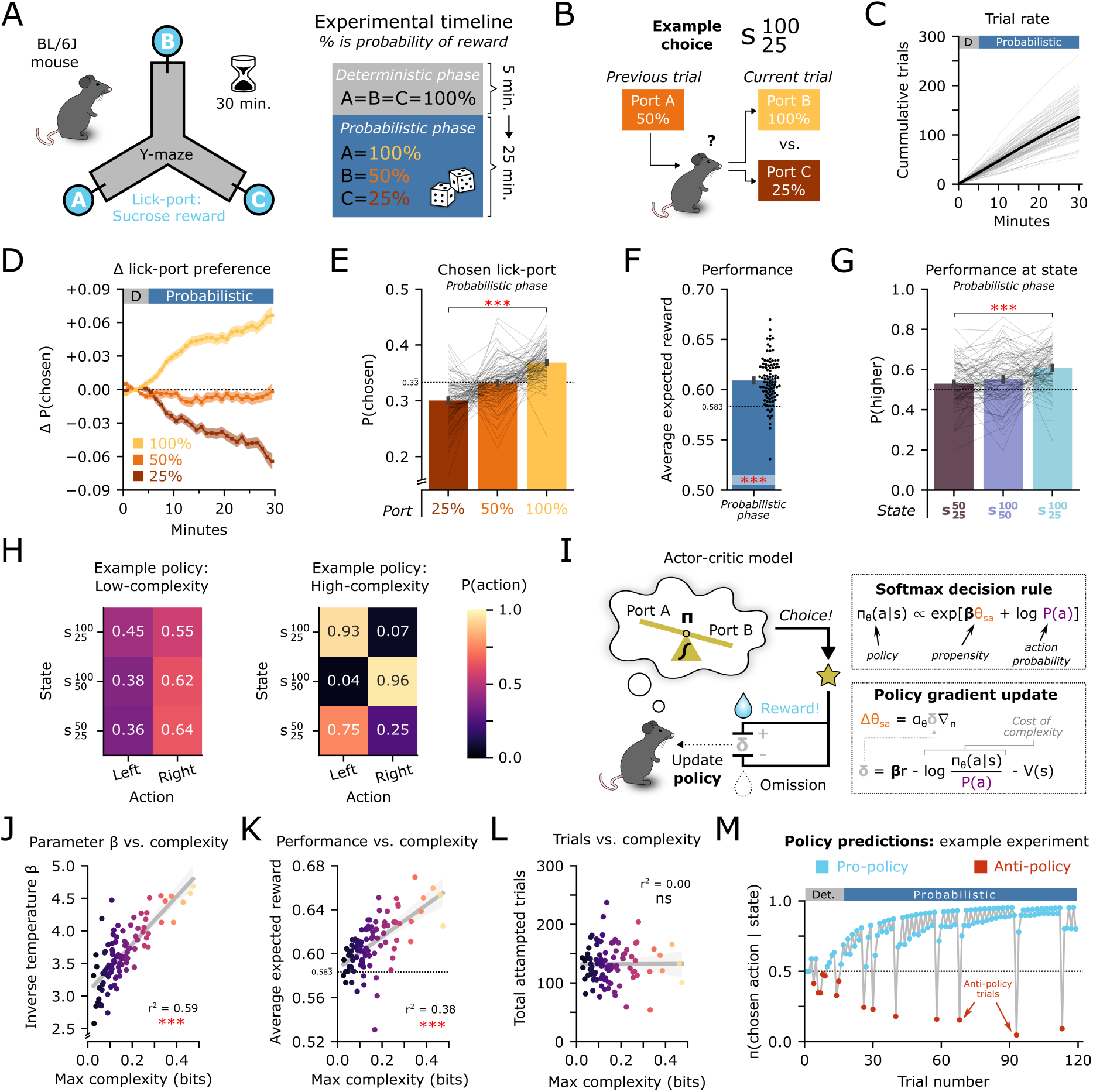
Experimental and modelling paradigm for quantifying decision policies in mice. **(A)** Schematic of the task and experimental timeline using n=97 mice. **(B)** Schematic of state-dependent decisions during the task. **(C)** Cumulative trials over time. **(D)** Lick-port preference over time baselined to the deterministic phase; smoothed by 5-trial moving average filter for visualization. **(E)** Probability to choose each lick-port during the probabilistic phase; rmANOVA. **(F)** Average reward expectation during the probabilistic phase. One-sample *t*-test against chance value of 0.583̄. **(G)** Probability to choose the higher-value lick-port at each state in the task; rmANOVA. **(H)** Example low and high complexity policies for state-dependent action selection. **(I)** Schematic of an actor-critic model consisting of a softmax decision rule and update via its policy gradient. See methods for more detail. **(J)** Maximum policy complexity versus model parameter β. **(K)** Maximum policy complexity versus performance in the probabilistic phase. **(L)** Maximum policy complexity versus total attempted trials in the probabilistic phase. **(M)** Example of model-predicted choices π(a|s) over trials. r^2^; coeXicient of determination. Error bars indicate SEM. *** *P* < 0.001, ns *P* > 0.05. Statistical details are provided in Supplementary Table 1.

Having determined that mice learned state-action contingencies in our task, we employed a computational model allowing us to study the complexity of their implemented policies. In this model, the hypothesis is that mice learn a policy *π* that maps states *s* to actions *a*. Recent work proposed an informational theoretical formalism where policies can be viewed as communication channels transmitting information about a current state (or context) to the probability of selecting an action in that state ^23 24^. Under the assumption that mice aim to maximize expected reward (by following policy *π*) while minimizing the costs of information processing, it can be shown the optimal policy is given by ^65^:

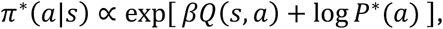

where *Q*(*s*, *a*) is the expected reward in state *s* after choosing a given action *a*, and *P*^∗^(*a*) is the marginal action probability. Crucially, parameter *β* determines the tradeoff between selecting the action with the highest reward and the marginal action probability, or in other words, how much information about the current state is considered versus a “default” action. Here, the mutual information *I*(*S*; *A*) between states and actions is defined as the *policy complexity* (methods). Given that mice must learn the state-action-reward contingencies via experience, we cast this optimization problem as a reinforcement learning formulation implemented through an actor-critic (AC) model, where mice are assumed to incrementally modify the policy based on reward feedback^24^ (Fig 1H-I, methods). A key hallmark of this formalism is that mice are penalized by the implementation of complex policies (Fig 1I). Based on leave-one-out cross-validation model comparison metrics, the AC policy compression model explained the data better than other competing models including standard versions of the RL learning model that do not consider policy complexity costs (Fig S1, methods).

We found that policy complexity increased with trial number (Fig S1), and that male and female mice attained similar levels of policy complexity (Fig. S1). The maximum achieved complexity at the end of each session was predictive of increased AC model parameter *β* (r^2^=0.59, *P*<0.001, Fig. 1J), and of higher overall task performance (r^2^=0.38, *P*<0.001, Fig. 1K). Interestingly, the maximum achieved policy complexity was not predictive of the total number of attempted trials (r^2^=0.00, *P*=0.947, Fig 1L). This may seem counterintuitive, as one expects that policy complexity will generally increase with experience. However, recall that the parameter *β* determines the overall capacity constraints, limiting the gains in policy complexity even after hundreds of trials. Together, these results may imply that mice aim at exerting a maximum metabolic investment determined by the internal states of the organism (this notion is studied in further detail in the causal manipulation experiments, below).

To gain more comprehensive insights into these observations, for some of the analyses in our subsequent imaging and causal manipulation experiments, we binarized trial-by-trial state action policies into two categories:

1. “Pro-policy”: *π*(*chosen action* | *state*) > 0.5), the “more likely” port predicted by the policy of the AC policy compression model.
2. “Anti-policy”: *π*(*chosen action* | *state*) < 0.5), the “less likely” port.

Anti-policy trials were less common, but still observed even after a mouse had over 100 trials of experience (Fig. 1M). One might interpret anti-policy decisions, which could be strategic exploratory-like decisions, as a failure of the AC model. However, our aim here is to exploit the interpretable predictions of this parsimonious model to question what processes would promote such apparent deviations (anti-policy) versus model policy alignment (pro-policy), which in turn reveal how adjustments in decision policy might be implemented at the neural level.

### Pre-choice HON dynamics encode decision policy

To measure HON activity during policy-driven action selection, we injected a viral vector carrying an orexin promotor-driven calcium sensor hORX.GCaMP6s into the lateral hypothalamus, and recorded the activity of n=102 HONs from four freely-behaving mice (Figure 2 A-B). To make sure that we exploit and understand the often-observed heterogeneity of HON responses ^53^, we first used a k-means clustering algorithm, based on which we identified at least four distinct subtypes of HON dynamics in the task (Fig. 2C). We found that neuron subtypes had a strong within-cluster correlation and they were heterogeneously distributed across mice (Fig. S2A-B).

**Fig. 2.**
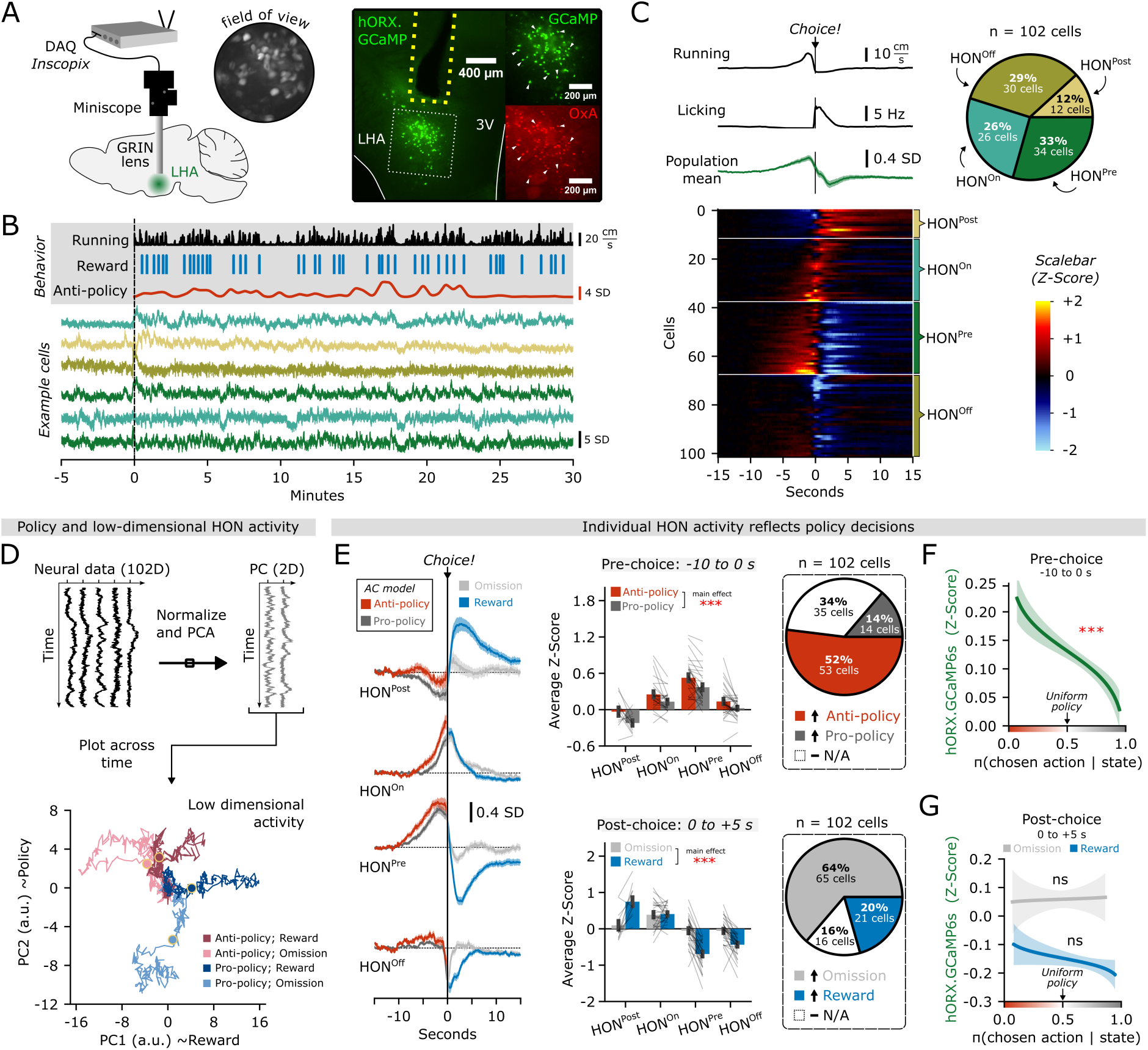
Hypothalamic neural signals correlate with policy decisions. **(A)** Schematic of miniscope recordings using hORX.GCaMP6s and representative histology from n = 4 implanted mice. The LHA region is magnified on the right for Orexin-A immunofluorescence. The white arrows indicate examples of overlapping fluorescence. 3V, third ventricle, LHA, lateral hypothalamus area. **(B)** Waterfall plot from an example experiment with photometry-aligned behavior and six example cells. **(C)** Choice-aligned running, licking, and photometry traces. Photometry is Z-Scored to a baseline -15 to -10 s before the choice. Population mean (green) is shown above 102 identified HONs, sorted by k-means clustering. The pie chart details the percentage abundance of each cluster. **(D)** Upper; schematic of low-dimensional principal components analysis. Lower; first and second principal components plotted across four trial-types. **(E)** Left; choice-aligned average photometry of each cluster color-coded by model predictions before the choice, and reward-outcome after the choice. Upper right; average photometry -10 to 0 s pre-choice separated by model predictions. Lower right; average photometry 0 to +5 s post-choice separated by reward outcome. Pie charts show the percentage of ROIs with higher (>0.1 SD) relative activity during a given event. **(F)** Linear regression to test if expit-transformed π(a|s) predicted the pre-choice photometry of pooled HONs. **(F)** Same as E, but for the post-choice epoch separated by reward-outcome. Error bars represent SEM. * *P* < 0.05, ** *P* < 0.01, *** *P* < 0.001, ns *P* > 0.05. Statistical details are provided in Supplementary Table 1.

Do HONs encode pre-choice policy and/or outcome events in the task? First, we computed a low-dimensional principal component (PC) representation of HON population dynamics and used the AC policy compression model policy predictions to visualize trial types in the low-dimensional space dynamics. The first and second PCs appeared to follow reward outcomes and decision policies respectively (Fig. 2D), suggesting that HON population dynamics carry information about the upcoming choice strategy and expected/received reward before the actual decisions are observed. A bulk-analysis of pre-choice HON activity similarly revealed significantly higher activity during anti-policy decisions (main effect of policy *F_1,98_* = 44.176, *P*<0.001, Fig. 2E) which was homogenous across subtypes (mixed ANOVA; *F_3,98_* = 0.211, *P*=0.889). Dynamics after a reward or omission were also pronounced, but heterogeneous across subtypes (mixed ANOVA; *F_3,98_* = 33.79, p<0.001) although most decreased their activity (main effect of reward *F_1,98_* = 33.739, *P*<0.001). Crucially, pre-choice HON activity was significantly negatively correlated with policy type before the actual choice was observed (*P*<0.001, Fig. 2F). However, HON activity during the feedback state (post-choice reward delivery or omission) was not related to policy, thus revealing a fundamental dissociation between actual (i.e. pre-choice) decision strategy and the post-choice decision consequences (Fig. 2G, Fig. S2C).

Is this relationship between policy and neural activity found in all neurons of the lateral hypothalamus? To answer this question, we conducted an identical set of analyses when imaging intermingled but non-overlapping hypothalamic neurons producing melanin-concentrating hormone (MCH)^66–68^. The activity of MCHs also displayed rapid and nonhomogeneous dynamics within the task, but only one activation subtype appeared to encode policy and reward (Fig. S3). This contrasts with the relation found for all HON activation cell subtypes (Fig 2E). Together, these data reveal that pre-choice HON population activity carries information about decision policy.

### HON activity modulates policy complexity depending on the organism’s nutritive state

It has been established that the overall level of HON activity is higher in energy-depleted states (e.g., nutrient deficit state) and lower in states of high energy reserves (e.g., nutrient surplus state)^29, 34–37^. In our previous imaging analyses (Fig. 2) mice were subject to a mild overnight (light cycle) food restriction before following-day experiments, thus suggesting relatively high levels of HON activity. The positive relationship between anti-policy decisions and HON activity (Fig. 2) and the prediction that “anti-policy” decisions lead to lower policy complexity (Fig. 3A) led us to hypothesize that, on the one hand, *fasted* mice whose HON activity would be transiently inhibited would adopt higher policy complexities. On the other hand, *fed* mice should not exhibit changes during inhibition given that HON activity should already be at low physiological levels.

**Fig. 3.**
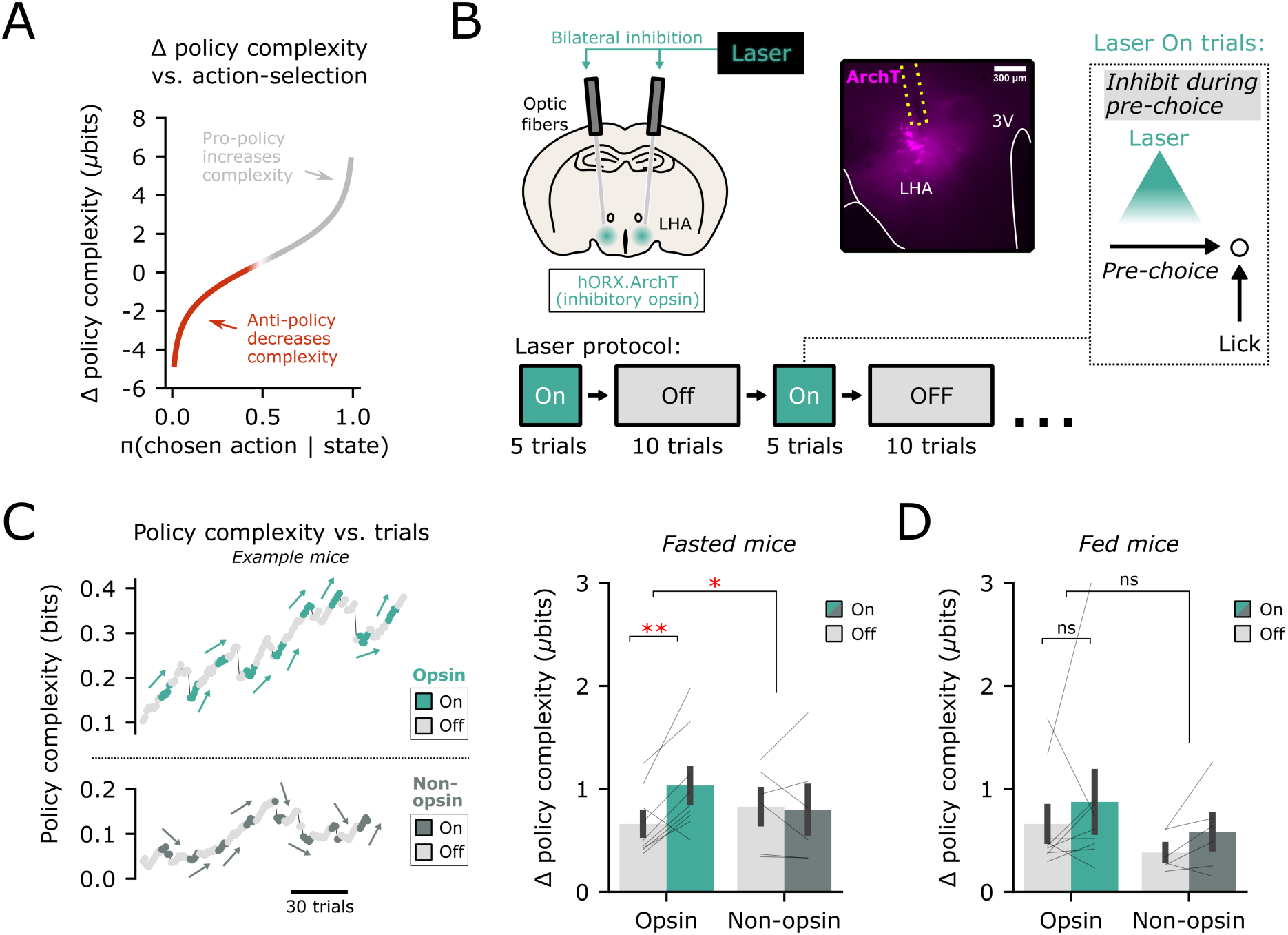
EGect of optosuppression of pre-choice HON activity on policy complexity. **(A)** Regression displaying how expit-transformed π(a|s) predicts the rate of change in policy complexity, using n = 97 mice from Figure 1. **(B)** Schematic showing inhibitory archaerhodopsin hORX.ArchT and representative histology of n = 9 mice. 3V, third ventricle, LHA, lateral hypothalamus area. Laser inhibition was applied 5 trials on, 10 trials oX, only during the pre-choice epoch. **(C)** Left; example trace of the rate of change in policy complexity in an opsin mouse and non-opsin control mouse. Right; rate of change in policy complexity during laser inhibition or sham trials in fasted opsin (n = 9) and non-opsin control (n = 6) mice. **(D)** Rate of change in policy complexity during laser inhibition or sham trials in fed mice. Error bars represent SEM. * *P* < 0.05, ** *P* < 0.01, *** *P* < 0.001, ns *P* > 0.05. Statistical details are provided in Supplementary Table 1.

To test this hypothesis, we suppressed pre-choice HON activity with temporally-targeted, HON-specific inhibitory optogenetics (inhibitory archaerhodopsin hORX.ArchT; Fig. 3B, methods) in blocks of 5 laser-on trials, interleaved with blocks of 10 laser-oI trials. This design allowed us to investigate whether the rate of change in policy complexity was higher during HON optosupression. In fasted mice, we found that HON optosupression led to an increase in the rate of change in policy complexity relative to laser-oI periods (paired t-test *P*=0.0075, Fig. 3C). This effect was significantly greater than control experiments using mice injected with a non-opsin fluorophore (mixed ANOVA *P*=0.042). However, in fed mice, we found the rate of change in policy complexity remained unaffected irrespective of intervention (*P*>0.05 for all tests, Fig. 3D). This result suggests HON signaling of nutrient state might be employed by downstream neural systems to regulate the amount of resources that should be invested to regulate information processing computations.

### Orexin signaling regulates policy complexity investment

While the optosupression experiment allowed us to establish that transient, pre-choice reductions in HON activity appear to enhance policy complexity at relatively high background levels of HON activation, these analyses did not allow us to establish how orexin signaling of nutrient reserves to downstream brain structures regulates policy complexity investment over longer temporal scales. HONs exert their downstream actions in part by releasing hypocretin/orexin peptide transmitters that act on brain-wide distributed hypocretin/orexin G-protein coupled receptors^69^. To investigate whether the hypocretin/orexin receptor activity regulates policy complexity, we reduced brain-wide hypocretin/orexin signaling with the dual hypocretin/orexin receptor antagonist almorexant^70^ (ALMO, methods). We hypothesized that ALMO should lead to an increased policy complexity investment. If this hypothesis holds, we should also observe that increased policy complexity due to ALMO should be reflected in lower reaction times and fewer attempted trials. For example, the more information needs to be processed on demixing input states and state-actions contingencies, the more processing time (i.e., pre-choice planning or deliberation) and post-choice consolidation time are required for planning and learning ^25, 71^.

Our initial observation was that mice injected with ALMO performed fewer trials than the same mice injected with vehicle (Fig. 4A). This effect was observed in both sexes and maintained over time (Fig. S5). Despite these strong effects on attempted trials, ALMO did not lead to a decrease in the final expected reward value (Fig. 4B and Fig S5), and did not change performance across states (Fig 4D). While initially counterintuitive, this immediately suggests that for a matched number of trials (from the session beginning), the suppression of hypocretin/orexin signaling should lead to the development of more complex policies (e.g., due to an increase in parameter *β* of the AC model, see Fig. 4E). In line with this reasoning, we found that ALMO increased estimates of the value of parameter *β* (within-subjects multiplicative effect *β*_&’()_=1.106, HDI_95%_ = [1.069, 1.148], *P*<0.001, Fig. 4E). Furthermore, ALMO resulted in the development of more complex policies at a similar number of trials from early into the session according to both the AC model (Fig. 4F) and using a model-free computation of policy complexity (Fig. 4G, methods).

**Fig. 4.**
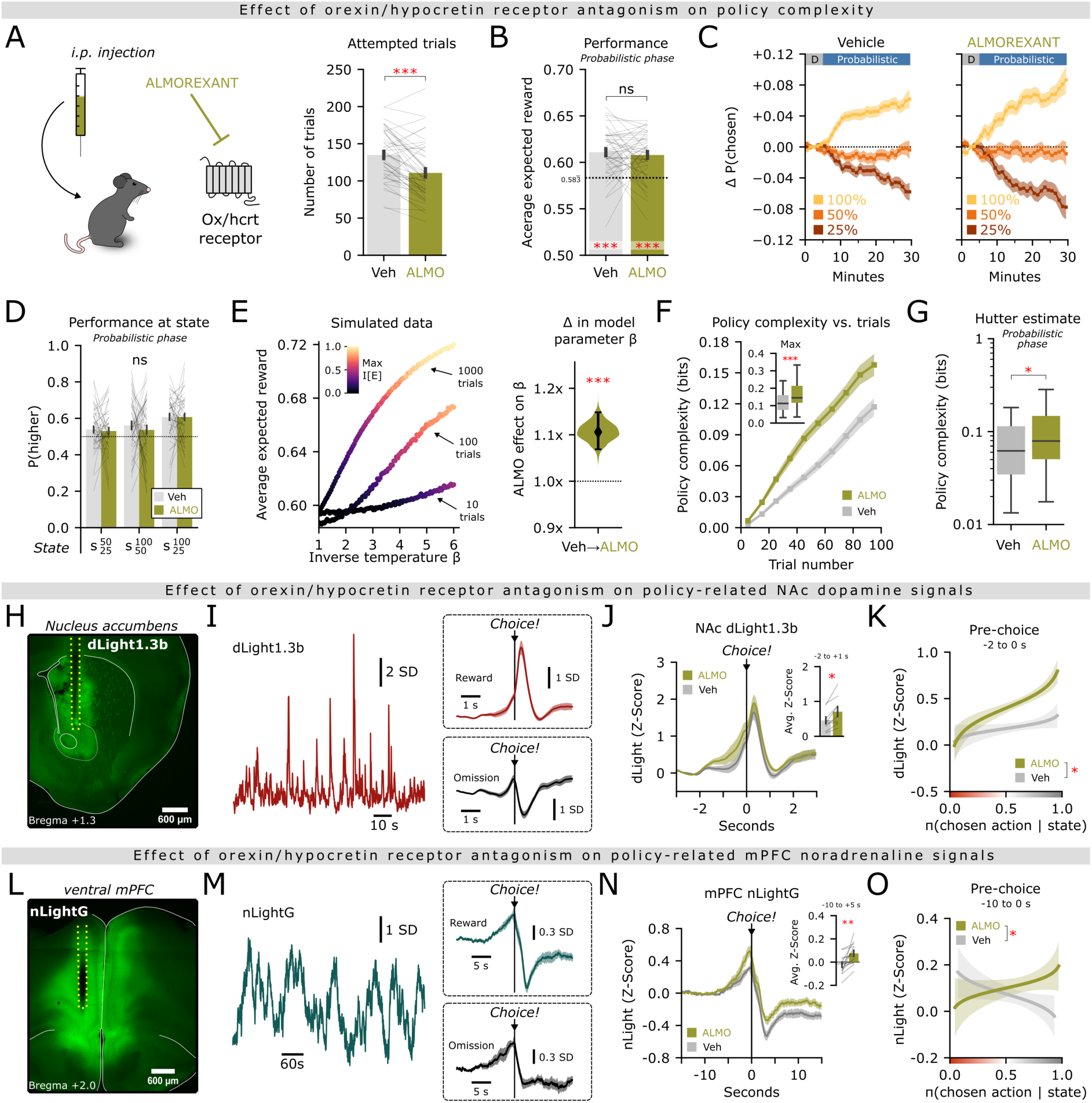
Control of policy complexity and related monoamine dynamics by hypocretin/orexin receptors. **(A)** Schematic showing injection of almorexant in n = 47 mice. Right; eXect of ALMO on total attempted trials. **(B)** EXect of ALMO on performance in the probabilistic phase. **(C)** Performance over time in both vehicle (left) and ALMO (right). **(D)** Probability to choose the higher-value lick-port at each state in the task; rmANOVA. **(E)** Left; simulated data plotting overall performance while increasing the value of β in the AC model. Color represents maximum policy complexity achieved. Right; posterior distribution of the eXect of ALMO on β from the AC model. Violin plots show median and highest density interval. **(F)** EXect of ALMO on the average policy complexity averaged across sessions. Inset shows mouse-averaged maximum policy complexity. Boxplots show median and inter-quartile range. **(G)** EXect of ALMO on Hutter estimate for policy complexity in the probabilistic phase. **(H)** Representative histology of N=9 mice expressing dLight1.3b in the NAc. **(I)** Example traces of dLight1.3b and reference signal. Right; average photometry traces of reward and omission trials. **(J)** Injection eXect on average dLight photometry of all trials Z-Scored to a baseline -3 to -2 s before the choice. Inset shows average of -2 to +1 s. **(K)** Linear model to test if expit-transformed π(a|s) predicted the pre-choice dLight photometry diXerently in vehicle or ALMO injected mice. Shaded region represents 95% CI. **(L)** Representative histology of N=14 mice expressing nLightG in the ventral mPFC. **(M)** Example traces of nLightG and reference signal. Right; average photometry traces of reward and omission trials. **(N)** Injection eXect on average nLightG photometry of all trials Z-Scored to a baseline -15 to -10 s before the choice. Inset shows average of -10 to +5 s. **(O)** Same as K, for nLightG. Error bars represent SEM. * *P* < 0.05, ** *P* < 0.01, *** *P* < 0.001, ns *P* > 0.05. Statistical details are provided in Supplementary Table 1.

Once again, we investigated whether these causal effects emerge due to a generalized action of the hypothalamus, or whether they are specific to HONs. We repeated the same set of experiments and analyses by applying an MCH-receptor antagonist (SNAP-94847, see methods) instead of ALMO. Despite a marginal increase in the number of attempted trials under SNAP-94847 relative to vehicle (Fig. S4), policy complexity remained unaltered at equivalent levels of experience.

### Aminergic neuromodulatory consequences of orexin signaling and decision policies

While the previous set of imaging analyses and causal manipulations strongly suggest that orexin signaling regulates policy complexity investment, there is ample evidence that the implementation of action plans and learning in tasks requiring learning under uncertainty is implemented by circuits such as the striatum and prefrontal cortex via dopaminergic^72^ and noradrenergic^73, 74^ neuromodulatory mechanisms, which are innervated by HON axons containing orexin/hypocretin transmitters^57–59^. Thus, we investigated whether ALMO affects the dynamics of noradrenaline and dopamine in medial prefrontal cortex (mPFC) and nucleus accumbens (NAc) respectively, and whether these possible downstream modulations influence the implementation of decision policies.

We recorded dopamine release in the NAc using fluorescent sensor dLight1.3b^75^. As expected, during the performance of the probability phase of our task, rewards induced an increase in dopamine activity, while omissions led to a reduction (Fig. 4H-I). Interestingly, peri-event dopamine responses were elevated by ALMO both before and after the decision (Fig 4J). Furthermore, under ALMO, the decision policy on a trial-by-trial basis became more predictive of pre-choice dopamine dynamics. That is, choices that led to anti-policy decisions enhanced pre-choice dopamine activity, while pro-policy trials attenuated dopamine activity in the NAc (Fig 4K).

We then turned to studying the influence of ALMO on prefrontal noradrenergic release. We recorded noradrenaline release in the ventral mPFC using a newly developed fluorescent sensor nLightG^76^ (Fig 4L). During our task, a pre-choice ramp-up in noradrenaline activity was followed by a rapid decrease following rewarded trials, while omissions saw a less pronounced reduction (Fig. 4M). While it is the first time that such mPFC noradrenaline dynamics are studied in this task, these results are similar to previously reported dynamics in noradrenaline-producing neurons of the locus coeruleus (LC) within the same time range^77^. Strikingly, levels of peri-event noradrenaline release were strongly elevated in a sustained manner under ALMO both before and after the decision (Fig. 4N). Furthermore, under ALMO, the decision policy at individual trials appeared to flip the contingency of pre-choice noradrenergic dynamics (Fig. 4O). That is, noradrenaline release had a positive slope for upcoming oI-policy decisions, which fits with the previously postulated role of noradrenergic release in promoting strategic exploratory shifts ^78^. The influence of ALMO showed similar modulations in a more dorsal region (the dorsal mPFC, Fig. S5), however, not as prominent as in the ventral area, and moreover the relation to decision policy was absent. This suggests a relative specificity of orexin signaling on noradrenaline release in the ventral mPFC (which could be considered homologous to the ventromedial prefrontal cortex in humans ^79^).

Once again we investigated the relative specificity of hypocretin/orexin receptors in policy control, finding that MCH receptor inhibitor (SNAP) did not lead to changes in dopamine or noradrenaline dynamics (Fig S4). Together, these data indicate that activation of hypocretin/orexin GPCRs influences policy-related dopamine and noradrenaline dynamics.

## DISCUSSION

Our results lead us to propose the following model. Nutritive deficiency activates HONs^29, 34–37^, which increases general locomotor arousal^56, 80–83^ and the number of attempts to obtain food (Fig. 4). At the same time, HON activity acts to reduce metabolically expensive^17, 84^ cognitive investment into each attempt, creating a less complex strategy in our task. HONs thus control the balance between cognitive vs physical effort. If it holds, this model suggests the existence of a general neurobiological mechanism for how resource deficiency states shift the organism from cognitive-dominant to movement-dominant strategies for maintaining performance.

Our theory suggests that HONs may be seated at the top of a hierarchical structure governing resource-dependent behavioral policies, given that they are well positioned to sense metabolic signals but also innervate brain areas that guide reward processing and learning updates while considering policy complexity costs^85^ (striatum, VTA), and action selection (basal ganglia). While other systems may take care of implementing cost-benefit analyses^30^ lower in the “hierarchy tree”, a missing piece in the puzzle has been how the brain signals the resource availability to control resource investment in the first place. This study places HONs as a critical missing link providing a more comprehensive mechanistic model of strategic learning accounting for the interplay between decision-making policies and metabolic/informational constraints. Beyond understanding the brain, this has implications for the design of AI systems, since it could inspire novel algorithms combining high-level resource monitors (analogous to HONs) with lower-level modules for cost-benefit analysis (akin to e.g. striatal dopamine systems), enabling systems to explicitly weigh the benefits of implementing complex behavioral policies against their related computational or energetic costs.

Our findings also have important implications for human health. Given that fasting regiments are increasingly popular as effective approaches for improving diverse aspects of health^86–88^, our findings suggest the possibility of maintaining certain complex cognitive operations in this context, for example through the use of hypocretin/orexin receptor antagonists already approved for human use^89–91^. Similarly, our findings also provide a HON-based hypothesis to be explored in future studies dissecting neurobiological causes of human diseases such as autism^92^ and schizophrenia ^93^, the latter of which has already been implicated in maladaptively reduced policy complexity.

## METHODS

### Animals

Female and male adult (>14 weeks old) wild-type C57BL/6J mice were housed in groups of up to five (same-sex) mice following a 12-h reversed light–dark cycle at 22 °C with 55% humidity. Animals were fed a diet of standard mouse chow (3430 Kliba Nafag). To ensure stable motivation and metabolic state, mice were subjected to a mild overnight (light cycle) food restriction before following day experiments, unless stated otherwise. Mice always had access to water. All behavioral tests were performed during the dark phase. Animal experiments followed Swiss Federal Food Safety and Veterinary Office Welfare Ordinance (Animal Welfare Ordinance 455.1, approved by the Zürich Cantonal Veterinary Office).

### Surgeries and viral vectors

For all mice requiring surgical operations, we first anesthetized mice with 5% isoflurane, which was lowered to 2% for the remainder of the surgery. Buprenorphine and site-specific lidocaine were used as additional analgesics. Injections and fiber / lens implantation were performed using a stereotaxic apparatus (Kopf Instruments) and a Nanoinject III injector. After surgery was completed, mice were given postoperative analgesia and monitored for recovery for at least two weeks before experiments began.

For microendoscope experiments, the activity of HONs (or MCH neurons) were recorded using a previously validated HON (or MCH) specific promoter-driven GCaMP6s sensor contained within an adeno-associated virus as previously described ^48, 49, 94–96^. For HON expression, we stereotaxically injected AAV1.hORX.GCaMP6s (2×10^13^ genome copies (GCs) per ml, Vigene Biosciences) unilaterally into the lateral hypothalamus. For MCH neurons, we similarly injected AAV9.pMCH.GCaMP6s (2×10^14^ GCs per ml, Vigene Biosciences). The coordinates for lateral hypothalamus were from bregma as follows: anteroposterior, −1.35; mediolateral, ±0.90; dorsoventral, −5.50, −5.35 and −5.10, 150 nl volume at 1 nl s^−1^ per site. The injection pipette was left in place after injection for at least 5 min to prevent reflux. Immediately after virus injection, a 27 gauge needle was attached to the stereotaxic holder and lowered to -5.00 dorsoventral at 50 µm/s, then removed at the same speed. A 0.6×7.3 ProView integrated GRIN lens (Inscopix) was lowered to -5.00 dorsoventral at 10 µm/s for the first 2 mm, and then 3.5 µm/s for the final 3 mm. Finally, the ProView baseplate was cemented in place to the skull.

For optogenetic experiments, we used the previously validated^80^ HON specific promoter-driven archaerhodopsin AAV1.hORX.ArchT.tdTomato (5 × 10^14^ GCs per ml, Vigene). We stereotaxically injected bilaterally into the lateral hypothalamus at the following coordinates from bregma: anteroposterior, −1.35; mediolateral, ±0.90; dorsoventral, −5.50, −5.35 and −5.10, 70-100 nl volume at 1 nl s^−1^ per site using a Nanoject III injector. The injection pipette was left in place after injection for at least 5 min to prevent reflux. For “control” experiments, we injected an adeno-associated virus containing non-opsin fluorophore tdTomato into HONs using AAV1.hORX.tdTomato (1.1 × 10^13^ GCs per ml, ETH Vector and Virus Production) at the same coordinates and volume. Optic fibers (200-μm diameter, 0.39 numerical aperture fiber with a 1.25-mm ceramic ferrule; Thorlabs) were implanted bilaterally above the injection site at -5.05 dorsoventral. Fibers were angled at ± 5° mediolateral to create space for bilateral stimulation and cemented in place to the skull.

For dopamine recording experiments, we used dLight1.3b sensor AAV9.CAG.dLight1.3b (7.9 × 10^12^ GCs per ml, UZH Viral Vector Facility). We stereotaxically injected unilaterally into the nucleus accumbens at the following coordinates from bregma: anteroposterior, +1.3; mediolateral, ±1.25; dorsoventral, −4.25, 300 nl volume at 1 nl s^−1^. The injection pipette was left in place after injection for at least 5 min to prevent reflux. For noradrenaline recording experiments, we used nLightG sensor AAV9.hSynapsin1.nLightG (1.4 × 10^13^ GCs per ml, UZH Viral Vector Facility). We stereotaxically injected into both the pre-limbic and infra-limbic mPFC at the following coordinates from bregma: pre-limbic; anteroposterior, +2.00; mediolateral, ±0.30; dorsoventral, −1.90, infra-limbic; anteroposterior, +2.00; mediolateral, ±0.30; dorsoventral,

−3.00. Both injections used a volume of 500 nL at 2 nl s^−1^. The injection pipette was left in place after injection for at least 10 min to prevent reflux. In all cases, optic fibers (200-μm diameter, 0.39 numerical aperture fiber with a 1.25-mm ceramic ferrule; Thorlabs) were implanted bilaterally 0.1 mm above the injection site and cemented in place to the skull. Pre-limbic fibers were angled at -20° anteroposterior to allow for simultaneous dual-site recording. We validated the nLightG sensor using a low dose of 10 mg kg^−1^ B.W. of desipramine, which extended nLightG dynamics (data not shown).

### Histology

To generate histological images, animals were first terminally anesthetized with pentobarbitone overdose and perfused intracardially using sterile PBS solution at pH 7.4, and then again using 4% paraformaldehyde solution in PBS. Brains were removed from the skulls and kept in 4 % paraformaldehyde overnight, at which point the solution was swapped with a 30% w/v sucrose solution (for dehydration) for another night. Brains were frozen in dry ice and stored at -80 °C until slicing using a vibrating microtome. Images were acquired using a fluorescence microscope (Eclipse Ti2, Nikon). When relevant, HONs were stained using goat anti-orexin-A (1:250 dilution, cat. no. sc-8070, Santa Cruz Biotechnology) and donkey anti-goat Alexa Fluor 546 (1:500 dilution, cat. no. A11056, Thermo Fisher Scientific).

### Probabilistic 3-armed bandit task

Experiments took place in a standard Y-Maze (Med Associates Inc.) with three identical arms forming an equilateral triangle. The end of each arm contained a custom-built liquid dispenser using a mechanical valve (cat. no. 161K011, NResearch) and a capacitor-based lick sensor (cat. no. AT42QT-1010, SparkFun Electronics) controlled via custom Python scripts and DAQ device (USB-6001, National Instruments). Rewards were calibrated to be 8 µL of 12% w/v sucrose water. An overhead infrared video camera allowed for remote monitoring of the task in dim / red light conditions; experimenters were not present in the room when behavior was recorded. The experiment duration was 30 minutes in which the mice could behave *ad libitum* in the maze.

When relevant, mice were first habituated to bilateral optic patch cable (Thorlabs) or microendoscope (Inscopix) tethering. Then, training and habituation to the Y-Maze took place over 4 days. During this time, the environment maintained a purely deterministic setting; every port had a 100% probability of dispensing a reward. Then, we introduced the experimental setting in which the 30 minute task occurred in two phases: the deterministic phase and probabilistic phase. The deterministic phase lasted from 0-5 minutes and was identical to the habituation environment (100% probability of reward). Then, the probabilistic phase began from 5-30 minutes in which each port was pseudo-randomly assigned one of three probabilities: 25%, 50%, or 100%. Critically, a reward could not be dispensed from the same port twice in a row, thus creating three unique states for the task. These contingencies were chosen based on previous designs ^97, 98^, which encouraged mice to engage in three unique probabilistic binary choices.

The experimental setting was repeated for several sessions, with the probabilities pseudorandomized per-mouse on every replicate. For most metrics in our analyses (i.e. number of trials, average performance), repeated measures across sessions were averaged. Chosen ports, individual licks, and their timestamps were recorded and saved to an HDF5 file, to be later analyzed using custom Python scripts. In postprocessing, we additionally aligned locomotor data recorded via an overhead infrared camera using a custom-trained deep network (ResNet18) trained to over 99% accuracy on over 5,000 manually labeled frames using PyTorch 2.2.0.

### Pharmacology and i.p. injections

The same animals received drug and vehicle in a cross-over design thus allowing for within-subjects analysis. Doses are as follows: A total of 30 mg kg^−1^ B.W. of the dual orexin/hypocretin receptor antagonist ALMO (Almorexant hydrochloride, MedChemExpress) or 20 mg kg^−1^ B.W. of the MCH receptor antagonist SNAP-94847 (cat. no. 3347 Torics Biosciences) or 10 mg kg^−1^ B.W. of the noradrenaline re-uptake inhibitor desipramine (cat. no. D3900, Sigma-Aldrich) was dissolved in 2% v/v dimethyl sulfoxide and potassium-buffered saline vehicle with 25% w/v 2-hydroxypropyl-β-cyclodextrin. The injection volume was 6 µL g^−1^ B.W. isovolumetric for drug and vehicle conditions. The concentration of drugs was chosen based on previous publications documenting their effects in mice^51, 96^. Drug and vehicle solutions were administered in a pseudorandom order via i.p. injections 40 minutes before performing behavior experiments. We habituated mice to i.p. injections of saline before the first experiment day.

### Microendoscope recordings and processing

Microendoscope (nVue System, Inscopix) recordings were performed using Inscopix acquisition software 2.1.0. The sampling rate was set to 10 Hz, exposure 99.9 ms, to capture a 640 x 400 video during the task. Max power was fixed as low as possible at ∼0.2 mW/mm^2^. Gain and focus were adjusted according to the Inscopix manual and kept constant per-mouse across recording sessions. Recordings of freely-moving animals took place first by allowing the mouse to habituate to the microendoscope headpiece over several days. Furthermore, during the six subsequent recording sessions the mice were given at least 10 minutes in an adjacent container to habituate to microendoscope attachment before being placed into the task.

We processed the data using Inscopix Data Processing Software 1.9.2. According to the user manual, preprocessing, spatial filter, motion correction, and ΔF/F were sequentially applied using default Inscopix settings. We used the cell identification method PCA/ICA with average cell diameter of 20 pixels, and the default settings over 1,000 maximum iterations. We took the conservative approach of only including cells that could be reliably identified in all six recording sessions using Inscopix’s longitudinal registration function. As an additional control, all ROIs were visually inspected based on shape and dynamics to verify accuracy. After alignment with behavioral data, we took the average activity of all HON ROIs to a rewarded trial ± 15 seconds, Z-Scored to a baseline of -15 to -10 seconds. We then used a k-means classifier (scikit-learn) to identify 3-4 clusters of activity patterns using ROIs as samples, and time as features. All ROIs were pooled for this analysis, and the classifier was blinded to the mouse-identity of each ROI.

ROI dynamics were aligned to two types of events: model prediction and trial outcome. Trial outcomes only depended on the binary reward status after a choice was made, while model predictions depended on the policy predictions *π*(*a*|*s*) of our AC model. For plotting and binned statistics in Figure 2, we used the increased thresholds of *π*(*a*|*s*) > 0.55 to characterize a pro-policy trial, and *π*(*a*|*s*) < 0.45 to characterize an anti-policy trial. In other words, the middle 10% of policy confidence was excluded to remove the trials which could not be distinctly binarized. However, when *π*(*a*|*s*) was used as a logit-transformed continuous metric (e.g. in the linear models) all trials were included.

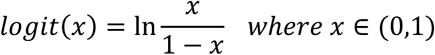

To generate low-dimensional PCA plots of neural population activity trajectories, we split all trials into four classifications separated by model prediction (pro/anti-policy) and trial outcome (reward/omission). We then calculated the average dynamics of each ROI across all classes, aligned to at trial ± 15 seconds, Z-Scored to a baseline of -15 to -10 seconds. To evaluate the component of activity non-selective to trial classification, we subtracted the average activity of each ROI calculated as a weighted-average across all trial types. Thereby, the resulting matrix contained features which represented how each ROI deviated from its average activity. The dynamics from the four trial-classes were then horizontally stacked, normalized, and passed into a PCA (scikit-learn). In this sense, the features (columns) represented 4×30 s of neural data, and samples (rows) represented individual ROIs. Finally, the principal components capturing the first and second most variance in the data were plotted against each other to form an activity trajectory for each trial classification.

### Optogenetics

For optogenetic inhibition, green laser (532 nm, Laserglow) stimulation was applied bilaterally to the lateral hypothalamus at ∼10 mW at fiber tip, laser continuously on with a 50 ms ramp on/oI. During the task, the laser was initially oI during the first 5 minutes (deterministic phase) and then repeated a 10-trial OFF (“Sham”) then 5-trial ON blocks during the probabilistic phase. Mice used in optogenetic experiments either expressed an opsin, or a fluorescent protein control. The experimenter was blinded to the cohort identity of each mouse. Repeated sessions (n = 3-5) were passed to an AC model. The parameter *β* was allowed to have a within-subjects effect of laser block, separated by cohort identity. Policy complexity predictions during the task were averaged and binned into laser ON/OFF blocks.

### Fiber photometry

Fiber photometry experiments were performed using a multifiber camera-based photometry system (Doric) using alternating illumination at 405 nm and 465 nm at 20 Hz, with a low average power of 20 µW at fiber tip. Emissions from dLight1.3b and nLightG were assumed to dominate the 465 nm wavelength. ΔF/F_0_ were generated by fitting the 465 nm emissions to the 405 nm using a first order least squares polynomial fit (numpy.polyfit). Peri-event signals were then taken at ± 3 seconds for dLight 1.3b and ± 15 seconds for nLight, determined by observing the natural dynamics of the sensors. Finally, z-scored traces were created by using a baseline of -3 to -2 seconds pre-choice (dLight1.3b) or -15 to -10 seconds pre-choice (nLightG). For visualization purposes only, photometry was smoothed using a 5-sample (250 ms) 1^st^ order Savitsky-Golay moving average filter; all statistics were performed on the raw data.

### Decision models

Because two consecutive rewards could not be obtained from the same port, behavior in the task can be modeled as a series of decisions between the two remaining available ports. The simplest model (bias model) assumed mice do not engage with the state-dependent nature of the task, and the action of going left or right was determined solely by *φ* ∈ (0,1), a fixed action probability.

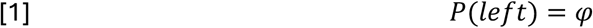

However, actions were likely state-dependent, forming a policy *π*(*a*|*s*) and these associations may be learned over time. We therefore implemented a classical reinforcement learning (RL) algorithm whereby a matrix Q dictates the expected reward for taking an action in each state. Thereby, the model described the probability of each action as a logistic softmax function, which is left implicit in subsequent equations).

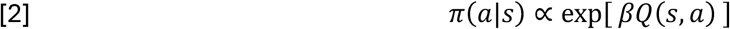

Within a softmax function, the *Q* matrix is multiplied by “inverse temperature” parameter which controls stochasticity governing the exploration-exploitation tradeoff. The 2×3 *Q* matrix was initialized to zeros on the first trial, and the Rescorla-Wagner model was used to update the *Q* matrix after each reward/omission. Specifically, each trial resulted in a binary outcome of *r* either a reward (1) or an omission (0) at trial *t*. Thereby the value of taking an action at a given state is updated via:

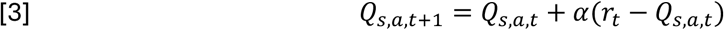

Where *⍺* ∈ (0,1) is the learning-rate parameter describing the rate at which value is updated with experience. The unchosen action was not updated. We refer to equations 2+3 as the RL model.

To formulate the actor-critic (AC) model in this paper, we assumed that the actor would depend on both state-dependent and marginal (state-independent) action probabilities, and that these two factors would directly interact such that the complexity of the policy would be subject to a capacity constraint. We therefore used a previously developed^23, 24^ capacity-constrained actor-critic model to explicitly update the policy with:

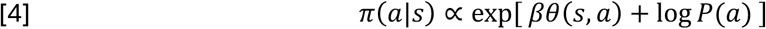

Where *P*(*a*) is the marginal action probability. We allowed marginal action probabilities to change with time, wherein the value *P*(*a*) was initialized 0.5 and updated every trial *t* via exponential moving average.

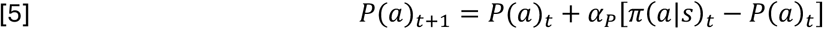

The parameter *⍺*_/_ ∈ (0,1) controls the rate at which marginal action probabilities are updated. *P*(*a*) was normalized by dividing by its sum on every timestep to ensure simplex conformity.

The matrix *θ* describes the propensity to select an action in a given state, which can be updated via a policy gradient algorithm:

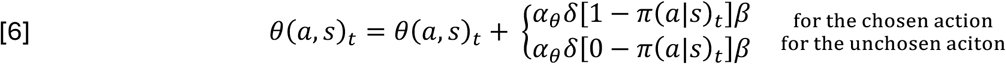

Where *⍺*_0_ ∈ (0,1) is the rate at which the actor’s policy gradient is updated, and *δ* is the prediction error of a critic *V^* (*s*):

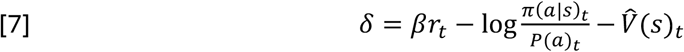

The critic was initialized to zeros and updated according to:

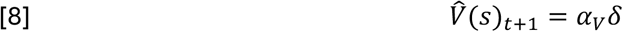

Where *⍺*_B_ ∈ (0,1) is the rate at which the critic is updated.

We also considered an extended version of the standard RL model, where we allowed the bias term to be dynamically updated following equation 5 (RLa model). However, note that this model does not incorporate policy complexity costs in the critic update rule. This allows us to study whether, in addition to marginal action probabilities influencing the choice policies, the actual policy costs are necessary to explain mice behavior.

To see an implementation of the models in Stan, line-by-line commented model code is available in this project’s online public repository.

A summary of the models used in this paper is given below:

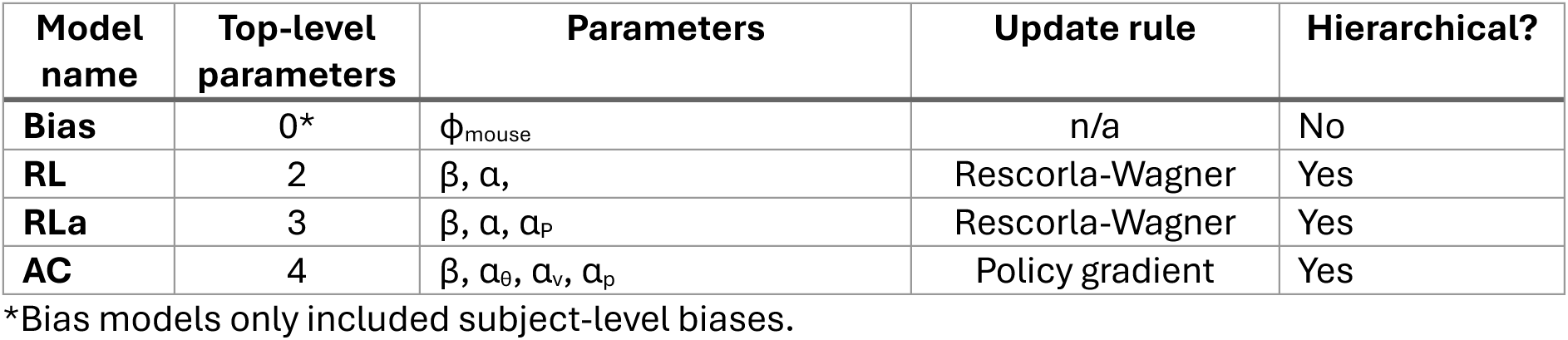

| Model name | Top-level parameters | Parameters | Update rule | Hierarchical? |
| --- | --- | --- | --- | --- |
| Bias | 0* | $\phi_{\text{mouse}}$ | n/a | No |
| RL | 2 | $\beta, \alpha,$ | Rescorla-Wagner | Yes |
| RLa | 3 | $\beta, \alpha, \alpha_p$ | Rescorla-Wagner | Yes |
| AC | 4 | $\beta, \alpha_\theta, \alpha_v, \alpha_p$ | Policy gradient | Yes |

\*Bias models only included subject-level biases.

### Model fitting and posterior statistics

Models RL, RLa, and AC were implemented as hierarchical Bayesian inference models, wherein individual parameters for each mouse were drawn from a population distribution. Subject-level parameters were drawn from population-level parameters using a non-centered parameterization. Lower-bounded parameters, (e.g. β) were constrained using an exponential link function, whereas dual-bounded parameters, (e.g. α) were constrained using an inverse logit link function. Subject-level effects were sampled using prior ∼*Normal*(0,1), population standard deviations (σ) ∼*Exponential*(1), and population means (µ) ∼*Logistic*(0,1). Note that in the case of dual-bounded parameters, the use of this logistic distribution and inverse-logit link function created a uniform prior.

The AC model was additionally parameterized to allow for a within-subjects design (e.g., in the ALMO experiments), when multiple conditions per subject existed. Thereby the drug effect, for example, could be modeled by a hierarchically-formulated parameter. Drug effects for lower-bounded parameters (i.e. β) were modeled using a multiplicative within-subjects effect which used the same bounds. Dual-bounded parameters (i.e. α_θ_) were modeled using an additive effect contained within the link function. For more specific details, line-by-line commented model code is available in this project’s online public repository.

Models were fit using Hamiltonian Monte Carlo sampling in Stan interfaced with CmdStanPy 1.2.4. For each model, we drew 2,500 warmup samples, followed by 2,500 real samples from each of four independent chains. A thinning of five was applied to the posterior chains, thus resulting in a set of 2,000 samples for each parameter. Split R-hat values were below 1.05 for all parameters, suggesting the chains converged to a similar posterior distribution. To compare models, we used the expected log pointwise predictive density leave-one-out cross-validation^99^ (ELPD-LOO) applied using the Python library ArviZ 0.14.0 (arviz.loo).

To determine significance of within-subject effects, the “*P* values” reported for these models are not frequentist *P* values, but instead directly quantify the probability of the reported effect differing from 0 (or 1 for multiplicative effects). These were computed using the posterior population distributions estimated for the parameter and represent the portion of the cumulative density functions that lies above/below 0 or 1 (depending on the direction of the effect). We did not assume directionality of any effects; all posterior “*P* values” were two-tailed. We reported effects as the median of the posterior distribution, alongside the 95% highest density interval (HDI) calculated using arviz.hdi.

### Deriving policy complexity

The complexity of a policy matrix can be quantified using the mutual information between its states and their associated action probabilities. In bits, the mutual information *I*^>^(*S*; *A*) can be calculated as:

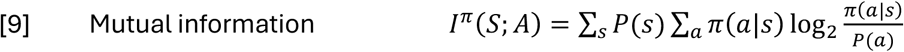

We input the average AC-model predicted *π*(*a*|*s*) matrix at each trial into equation 10, where *P*(*a*) is the marginal action probability and *P*(*s*) is the probability of being in a given state (calculated using the transition matrix and a system of linear equations). For statistical comparison purposes, we took the maximum mutual information attained within the first 100 trials averaged per-mouse. For visualization only, we trial-aligned each individual session and averaged within 10-trial bins. Finally, to approximate the slope policy complexity (Fig. 3) we took the calculated mutual information vector (length = trials) and differentiated using a Savitzky-Golay filter of length 5 and order 1. The Hutter approximation ^100^ was used to generate an alternative quantification of policy complexity that did not depend on the output of an AC model. From each session, we took all trials from the probabilistic phase and calculated the Hutter estimator between states (*s*) and actions (*a*). First, a co-occurrence matrix *N* was initialized with a prior *τ*.

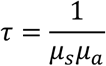

Where *μ*_*_and *μ*, are the number of unique states and actions. The policy complexity was calculated using the digamma function-based correction:

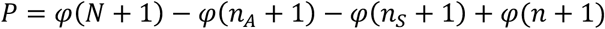

where *φ*(·) is the digamma function, *n*_&_and *n*_C_ are the marginal counts of *N*, and *n* is the total count. The final policy complexity *I*(*S*; *A*) is defined as:

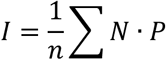

### Statistics

Behavioral data was acquired and processed using python version 3.9-12. Frequentist statistics were performed using python via SciPy, statsmodels, and Pingouin libraries. Broadly, we did not exclude any animals or recording sessions based on any performance criteria of their behavior. Regrettably, a small number of sessions had to be excluded due to software failures, mechanical failures of lick-ports, or restricted mouse movement due to optic patch cable tangling. A small number of mice also had to be excluded from photometry analysis due to no obvious dynamics.

Most data were represented as a mean and standard error on the mean (SEM). In a few metrics, the data was clearly not normally distributed, in which case nonparametric tests were used alongside box and whisker plots. Box properties represented the median and quartiles. For plotting purposes, whiskers extended to the maximum and minimum of the distribution after excluding outliers outside 1.5 × the inter-quartile range. Outlier exclusion was for visualization only; nonparametric tests included all data. Regardless of the test, significance was defined as the following *P* values: \**P* < 0.05, \*\**P* < 0.01, \*\*\**P* < 0.001, NS represents *P* > 0.05. In the case where multiple conditions were simultaneously compared, we used the Bonferroni correction to generate more conservative *P*-values. For linear modeling, predicted mean and 95% confidence intervals are displayed as a shaded patch, when relevant. We made no directional assumptions with any of our statistical tests; all tests were two-tailed. Statistical information for all figures is summarized in supplementary table 1.

## Data availability

Data will be deposited in a publicly available database (osf.io) before time of publication.

## Acknowledgements

This work was funded by ETH Zürich. DB, RP, and ALT conceived the study and designed the protocol, with contributions from DPR and NG. ALT and EFB performed the surgeries. ALT, CDP, and DG performed the experiments. TP provided and advised on the dopamine/noradrenaline sensors. DB, RP, and ALT wrote the text with inputs from NG. The authors have no competing interests to declare

---

**Supplementary Table 1.**

| Fig | Test | Statistic | p-value | Sample size |
| --- | --- | --- | --- | --- |
| **Fig. 1** |  |  |  |  |
| 1E | rmANOVA | $F_{2,192} = 51.86$ | $P < 0.001$ | n = 97 |
| 1F | 1 sample $t$-test | $t_{96} = 11.165$ | $P < 0.001$ | n = 97 |
| 1G | rmANOVA | $F_{2,192} = 11.98$ | $P < 0.001$ | n = 97 |
| 1J | OLS regression | $t = 11.802$ | $P < 0.001$ | n = 97 |
| 1K | OLS regression | $t = 7.588$ | $P < 0.001$ | n = 97 |
| 1L | OLS regression | $t = 0.067$ | $P = 0.947$ | n = 97 |
| **Fig. 2** |  |  |  |  |
| 2E | rmANOVA (policy -10 to 0 s) | $F_{3,98} = 0.479$ | $P = 0.698$ | n = 102 |
| 2E | rmANOVA (reward 0 to +5 s) | $F_{3,98} = 33.79$ | $P < 0.001$ | n = 102 |
| 2F | OLS regression | $t = -5.443$ | $P < 0.001$ | n = 46,881 |
| 2G | OLS regression (omission) | $t = -0.168$ | $P_{\text{adj}} = 1.000$ | n = 14,400 |
| | OLS regression (reward) | $t = -1.780$ | $P_{\text{adj}} = 0.150$ | n = 32,481 |
| **Fig. 3** |  |  |  |  |
| 3C | Paired $t$-test | $t_8 = 3.518$ | $P = 0.008$ | n = 9 |
| | Mixed ANOVA | $F_{1,13} = 5.960$ | $P = 0.030$ | - |
| 3D | Paired $t$-test | $t_8 = 0.883$ | $P = 0.402$ | n = 9 |
| | Mixed ANOVA | $F_{1,13} = 0.002$ | $P = 0.969$ | - |
| **Fig. 4** |  |  |  |  |
| 4A | Paired $t$-test | $t_{46} = 5.847$ | $P < 0.001$ | n = 47 |
| 4B | Paired $t$-test | $t_{46} = 0.614$ | $P = 0.542$ | n = 47 |
| | 1 sample $t$-test (vehicle) | $t_{46} = 7.331$ | $P_{\text{adj}} < 0.001$ | n = 47 |
| | 1 sample $t$-test (almorexant) | $t_{46} = 6.810$ | $P_{\text{adj}} < 0.001$ | n = 47 |
| 4D | Paired $t$-test | $F_{2,92} = 0.134$ | $P = 0.874$ | n = 47 |
| 4E | Bayesian posterior | HDI<sub>95%</sub> [1.069, 1.148] | $P < 0.001$ | 2,000 draws |
| 4F | Wilcoxon signed-rank test | $z = 234.0$ | $P < 0.001$ | n = 47 |
| 4G | Wilcoxon signed-rank test | $z = 370.0$ | $P = 0.040$ | n = 47 |
| 4J | Paired $t$-test | $t_8 = 3.628$ | $P = 0.013$ | n = 9 |
| 4K | OLS regression (interaction) | $t = 2.387$ | $P = 0.017$ | n = 7,489 |
| 4N | Paired $t$-test | $t_{12} = 3.703$ | $P = 0.006$ | n = 13 |
| 4O | OLS regression (interaction) | $t = 2.474$ | $P = 0.013$ | n = 7,665 |
| **Sup. 1** |  |  |  |  |
| S1D | Unpaired $t$-test | $t = 3.558$ | $P = 0.001$ | n = 42♂, 55♀ |
| S1E | Unpaired $t$-test | $t = 1.706$ | $P = 0.091$ | n = 42♂, 55♀ |
| | 1 sample $t$-test (male) | $t_{41} = 9.053$ | $P_{\text{adj}} < 0.001$ | n = 42 |
| | 1 sample $t$-test (female) | $t_{54} = 7.165$ | $P_{\text{adj}} < 0.001$ | n = 55 |
| S1F | Mann–Whitney $U$ test | $U = 1353.0$ | $P = 0.150$ | n = 42♂, 55♀ |
| **Sup. 2** |  |  |  |  |
| S2C | OLS regression | $t = -1.596$ | $P < 0.114$ | n = 102 |
| **Sup. 3** |  |  |  |  |
| S3D | rmANOVA (interaction effect) | $F_{2,40} = 8.143$ | $P < 0.001$ | n = 43 |
| | rmANOVA policy (main effect) | $F_{1,40} = 8.143$ | $P = 0.003$ | n = 43 |
| | Paired $t$-test (MCH<sup>Pre</sup>) | $t_{14} = 3.612$ | $P = 0.003$ | n = 15 |
| S3E | rmANOVA (interaction effect) | $F_{2,40} = 14.826$ | $P < 0.001$ | n = 43 |
| | rmANOVA reward (main effect) | $F_{1,40} = 19.948$ | $P < 0.001$ | n = 43 |
| | Paired $t$-test (MCH<sup>Pre</sup>) | $t_{14} = 7.296$ | $P < 0.001$ | n = 15 |
| **Sup. 4** |  |  |  |  |
| **S4A** | Paired $t$-test | $t_{46} = 2.607$ | $P = 0.012$ | n = 47 |
| **S4B** | Paired $t$-test | $t_{46} = 0.265$ | $P = 0.012$ | n = 47 |
| | 1 sample $t$-test (vehicle) | $t_{46} = 7.331$ | $P_{\text{adj}} < 0.001$ | n = 47 |
| | 1 sample $t$-test (SNAP) | $t_{46} = 6.823$ | $P_{\text{adj}} < 0.001$ | n = 47 |
| **S4C** | rmANOVA |  |  | n = 47 |
| **S4D** | Bayesian posterior test | HDI<sub>95%</sub> [0.963, 1.050] | $P = 0.784$ | 2,000 draws |
| | Wilcoxon signed-rank test | $z = 527.0$ | $P = 0.702$ | n = 47 |
| **S4E** | Paired $t$-test | $t_8 = 0.406$ | $P = 0.696$ | n = 9 |
| **S4F** | Paired $t$-test | $t_{12} = 0.302$ | $P = 0.768$ | n = 13 |
| **Sup. 5** |  |  |  |  |
| **S5A** | Paired $t$-test (male) | $t_{24} = 3.722$ | $P_{\text{adj}} = 0.002$ | n = 25 |
| | Paired $t$-test (female) | $t_{21} = 3.739$ | $P_{\text{adj}} < 0.001$ | n = 22 |
| **S5B** | Paired $t$-test (male) | $t_{24} = 1.155$ | $P_{\text{adj}} = 0.259$ | n = 25 |
| | Paired $t$-test (female) | $t_{21} = 0.148$ | $P_{\text{adj}} = 0.884$ | n = 22 |
| **S5C** | Paired $t$-test (25%) | $t_{47} = 0.443$ | $P_{\text{adj}} = 1.000$ | n = 47 |
| | Paired $t$-test (50%) | $t_{47} = 0.077$ | $P_{\text{adj}} = 1.000$ | n = 47 |
| | Paired $t$-test (100%) | $t_{47} = 0.503$ | $P_{\text{adj}} = 1.000$ | n = 47 |
| **S5E** | Bayesian posterior test | HDI<sub>95%</sub> [0.0004, 0.0017] | $P < 0.001$ | 2,000 draws |
| | Wilcoxon signed-rank test | $z = 273.0$ | $P = 0.002$ | n = 47 |
| **S5F** | Paired $t$-test | $t_{12} = 4.316$ | $P = 0.002$ | n = 13 |
| | OLS regression (interaction) | $t = 1.167$ | $P = 0.243$ | n = 7,665 |

$P_{\text{adj}}$ implies a Bonferroni correction for multiple comparisons
We make no assumptions of directionality; all tests are two-tailed.

**Fig. S1.**
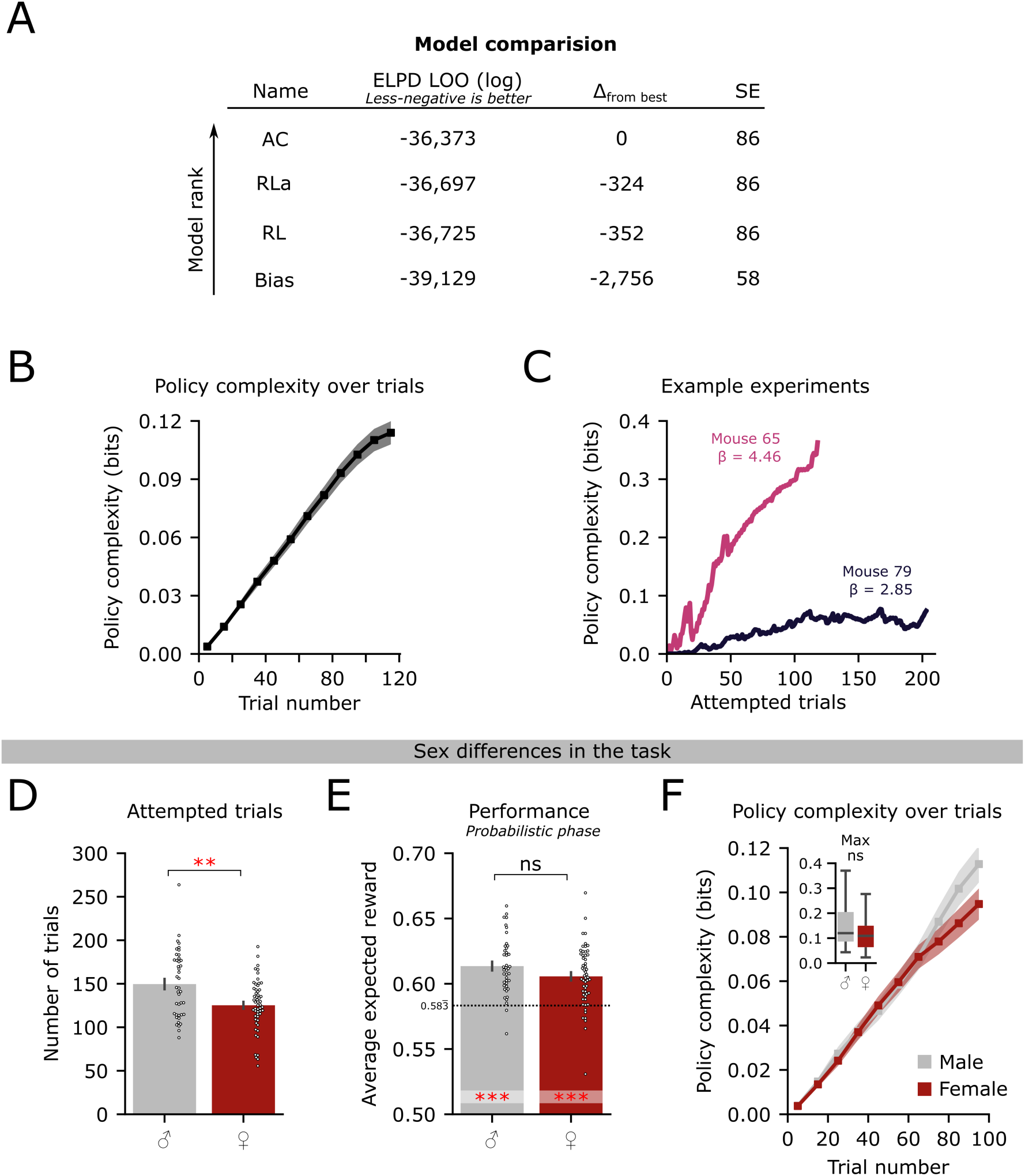
Model characterization and sex diGerences. **(A)** Model comparison of four models fit to the same dataset from Figure 1. **(B)** Average policy complexity over trials from all sessions in the task. Shaded region represents SEM. **(C)** Example sessions in which a high-β mouse had a more complex policy than a low-β mouse despite attempting fewer trials. **(D)** EXect of sex on the total number of attempted trials. **(E)** EXect of sex on performance during the probabilistic phase. **(F)** EXect of sex on the average policy complexity averaged across sessions. Inset shows mouse-averaged maximum policy complexity. Error bars and shaded regions represent SEM. Box plots show median and IQR. *** *P* < 0.001, ns *P* > 0.05. Statistical details are provided in Supplementary Table 1.

**Fig. S2.**
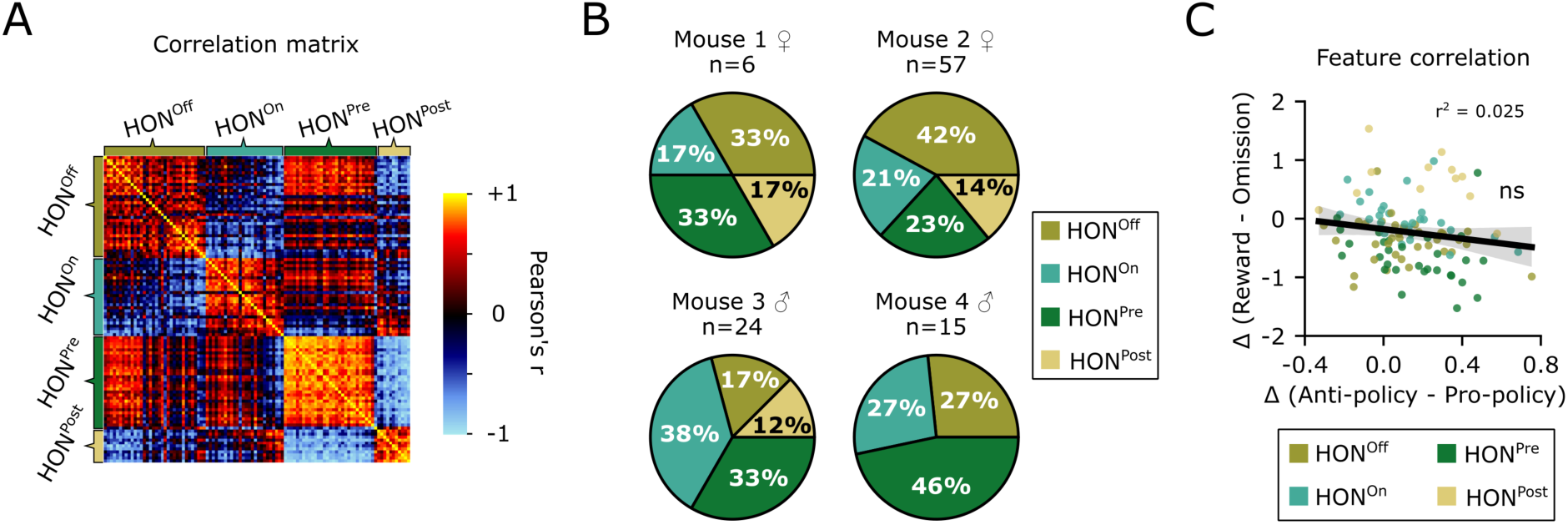
Additional characterization of HON dynamics. **(A)** Within and between cluster correlation of all HONs aligned to a choice ± 15 s. **(B)** Proportion of identified subtypes across all recorded mice. **(C)** Correlation of individual cell responses to both policy decisions and reward-outcome. Shaded region represents 95% CI. ns *P* > 0.05, Statistical details are provided in Supplementary Table 1.

**Fig. S3.**
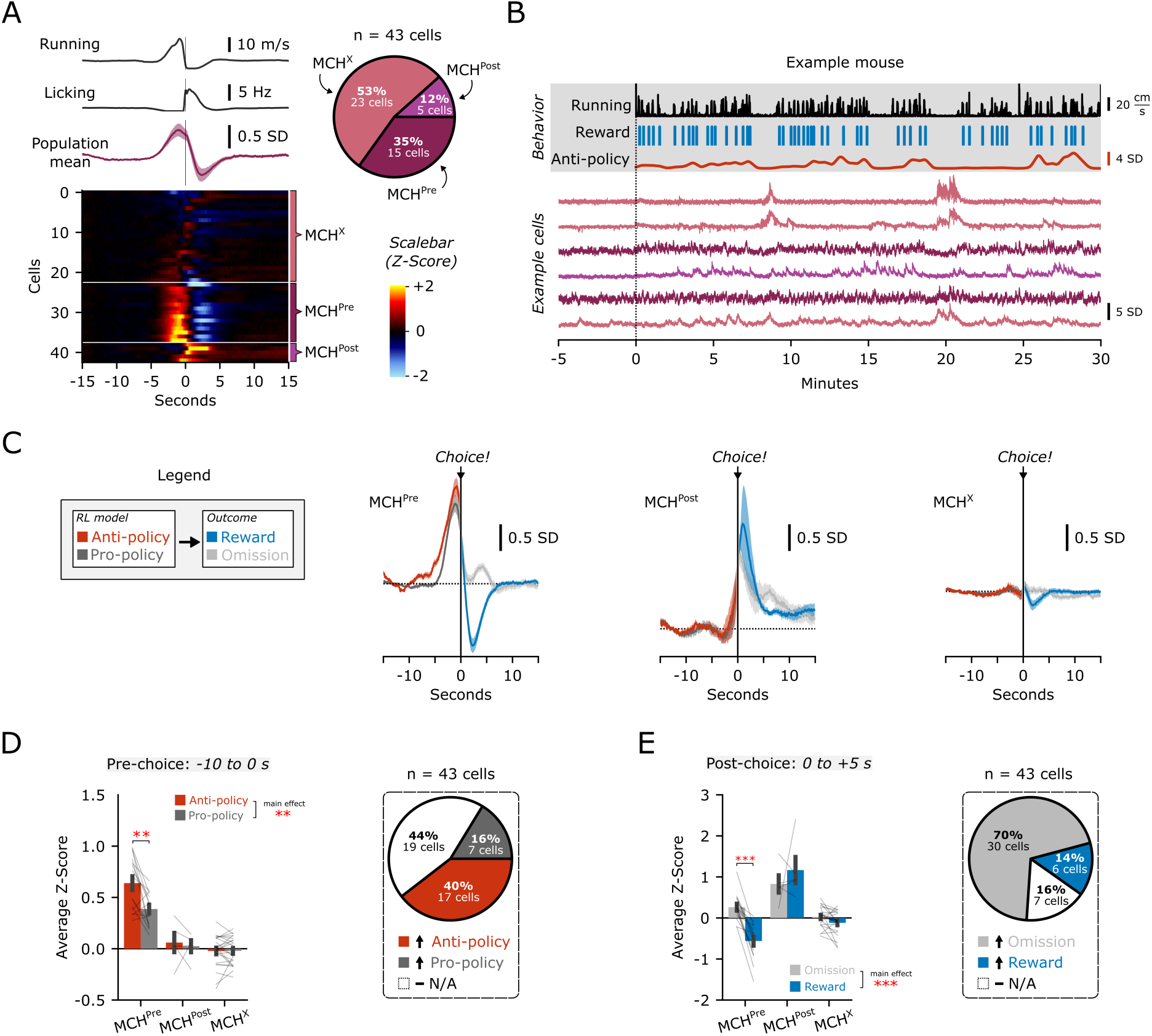
MCH neuron activity in the bandit task. **(A)** Choice-aligned running, licking, and photometry traces. Photometry is Z-Scored to a baseline -15 to -10 s before the choice. Population mean (purple) is shown above 43 identified MCH neurons from n = 5 mice, sorted by k-means clustering. The pie chart details the percentage abundance of each cluster. **(B)** Waterfall plot from an example experiment with photometry-aligned behavior and six example cells. **(C)** Choice-aligned average photometry of each cluster color-coded by model predictions before the choice, and reward-outcome after the choice. **(D)** Average photometry -10 to 0 s pre-choice separated by model predictions. Pie charts show the percentage of ROIs with higher (>0.1 SD) relative activity during a given event **(E)** Average photometry 0 to +5 s post-choice separated by reward outcome. Pie charts show the percentage of ROIs with higher (>0.1 SD) relative activity during a given event Error bars represent SEM. * *P* < 0.05, ** *P* < 0.01, *** *P* < 0.001, ns *P* > 0.05. Statistical details are provided in Supplementary Table 1.

**Fig. S4.**
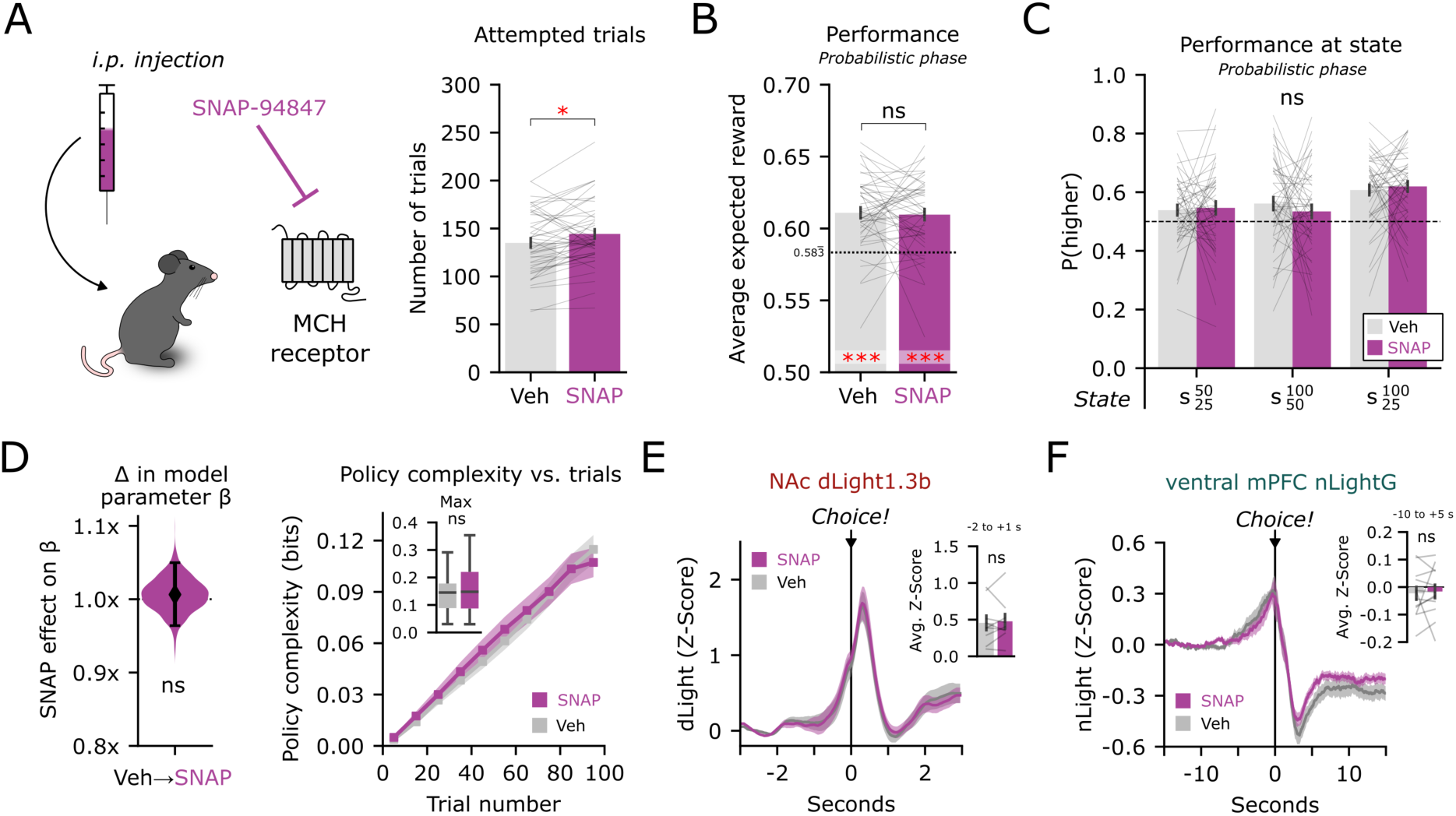
Antagonism of MCH receptors in the bandit task. **(A)** Schematic showing injection of SNAP-94847. Right; eXect of SNAP on total attempted trials. **(B)** EXect of SNAP on performance in the probabilistic phase. **(C)** Probability to choose the higher-value lick-port at each state in the task; rmANOVA. **(D)** Left; posterior distribution of the eXect of ALMO on β from the AC model. Violin plots show median and highest density interval. Right; eXect of SNAP on the average policy complexity averaged across sessions. Inset shows mouse-averaged maximum policy complexity. Boxplots show median and inter-quartile range. **(E)** Injection eXect on average photometry of all trials Z-Scored to a baseline -3 to -2 s before the choice. Inset shows average of -2 to +1 s. **(F)** Injection eXect on average photometry of all trials Z-Scored to a baseline -15 to -10 s before the choice. Inset shows average of -10 to +5 s. Error bars represent SEM. * *P* < 0.05, ns *P* > 0.05. Statistical details are provided in Supplementary Table 1.

**Fig. S5.**
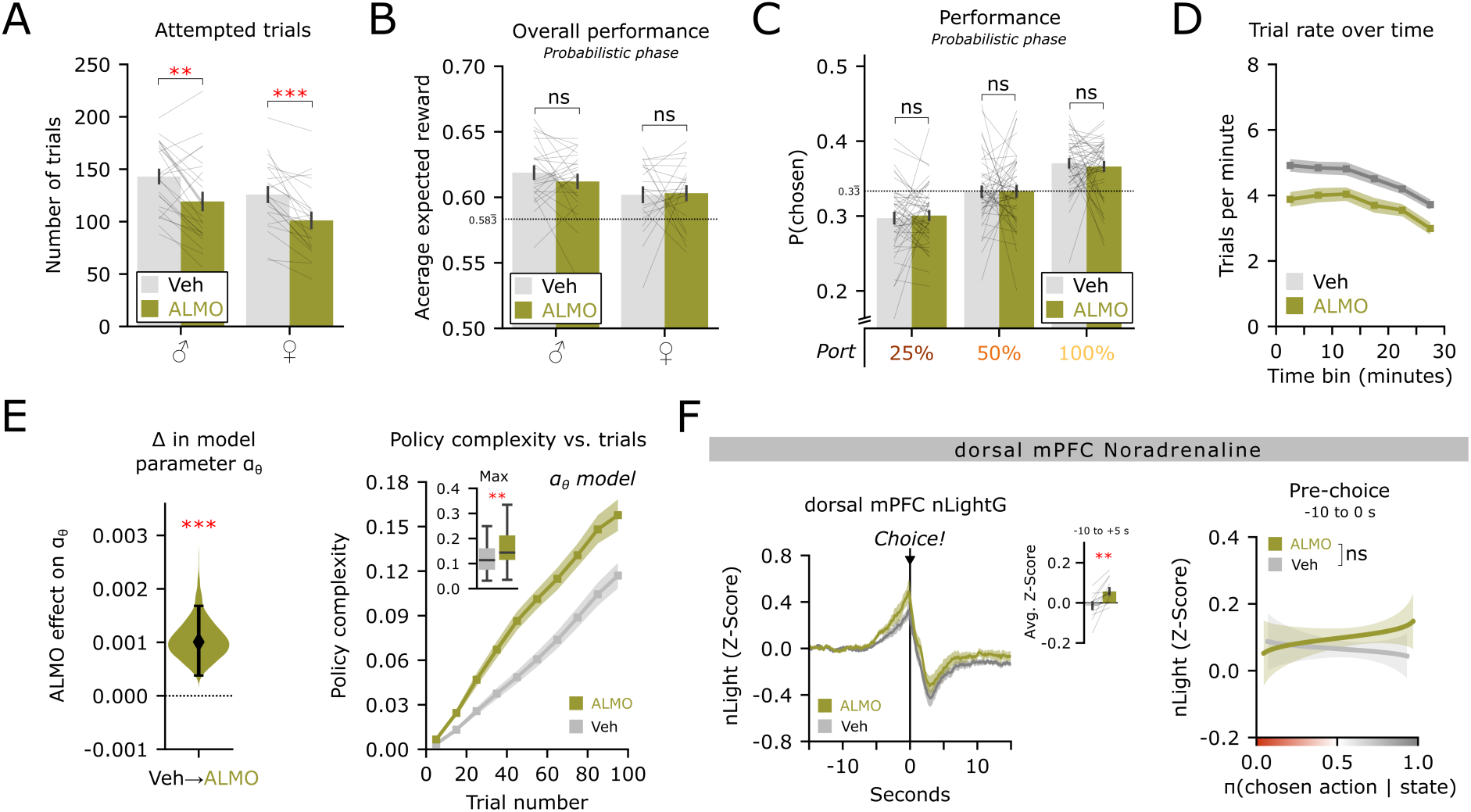
Additional characterization of ALMO eGects. **(A)** EXect of ALMO injection on total attempted trials by sex. **(B)** EXect of ALMO on performance in the probabilistic phase by sex. **(C)** EXect of ALMO probability to choose each lick-port during the probabilistic phase. **(D)** EXect of ALMO on trial rate in five minute bins. **(E)** Left; Posterior distribution of the eXect of ALMO on an AC model where the eXect of ALMO was on the αθ parameter. Violin plot shows the median and highest density interval. Right; EXect of ALMO on the average policy complexity averaged across sessions. Inset shows mouse-averaged maximum policy complexity. Boxplots show median and inter-quartile range. **(F)** Left; ALMO eXect on average nLightG photometry from the dorsal mPFC. Inset shows average of -10 to +5 s. Right; Linear model to test if expit-transformed π(a|s) predicted the pre-choice photometry diXerently in vehicle or ALMO injected mice. Shaded region represents 95% CI. Error bars represent SEM. ** *P* < 0.01, *** *P* < 0.001, ns *P* > 0.05. Statistical details are provided in Supplementary Table 1.

